# Metagenomic insights into dose-dependent alterations in the tobacco rhizosphere microbiome by pyrroloquinoline quinone

**DOI:** 10.64898/2026.10.08.757804

**Authors:** Liwei Liu, Yanqi Liang, Jiahao Cai, Yan Cheng, Chang An, Chao Shi, Chongrong Ke, Yuan Qin, Chunmei Lai

## Abstract

Pyrroloquinoline quinone (PQQ) is known to promote plant growth, yet how the rhizosphere microbiome varies with PQQ concentration remains poorly understood. We applied metagenomic sequencing to characterize the taxonomic composition and functional potential of the tobacco rhizosphere microbiome across PQQ treatments at 0.5, 1 and 2 μmol/L plus an untreated control. The rhizosphere microbiome was dominated by Pseudomonadota and Actinomycetota and differed among treatments in a concentration-associated, non-monotonic manner. The most pronounced differences were observed at 1 μmol/L, where alpha diversity showed dimension-specific variation and beta-diversity analysis resolved the most distinct community profile. Taxonomic and functional profiles varied concurrently, with *Streptomyces*, *Sphingomonas*, *Rhodanobacter* and *Lysobacter*, together with functions related to biosynthesis, energy metabolism, colonization and environmental sensing, showing higher relative abundance at this concentration. No broad-spectrum antimicrobial effect was detected at the community level, and antibiotic resistance gene abundance showed no significant differences among treatments after correction for multiple testing. These findings provide a basis for further evaluation of PQQ in modulating the rhizosphere microbiome in continuous cropping systems.

**Statement of significance:** For the first time in tobacco, we demonstrate that PQQ governs the rhizosphere microbiome in a concentration-associated yet non-monotonic manner, resolving the most distinctive community architecture at 1 μmol/L, where Pseudomonadota and Actinomycetota predominate and alpha diversity shifts dimension-specifically. Notably, taxonomic and functional trajectories co-vary, driving concurrent enrichment of *Streptomyces*, *Sphingomonas*, *Rhodanobacter* and *Lysobacter* together with functions underpinning biosynthesis, energy metabolism, colonization and environmental sensing. Crucially, the absence of broad-spectrum antimicrobial activity and of significant antibiotic resistance gene changes positions PQQ as a novel, microbiome-targeted candidate, supporting its further evaluation in modulating rhizosphere microbiomes in continuous cropping systems.

## 1. Introduction

The rhizosphere serves as a pivotal interface connecting plant roots, soil matrices, and microbial assemblages, governing the translocation of nutrients, water, and bioactive substances within root systems [1]. Rhizosphere microbial communities not only mediate soil nutrient activation and contaminant degradation, but also modulate plant physiological performance by regulating hormonal homeostasis and secondary metabolism [2]. Such plant–microbe interactions are shaped by bilateral selective filtering. Edaphic factors, including soil pH and organic matter, structure microbial community assembly, while plant root exudates reciprocally reshape microbiota composition and functional profiles [3,4]. This bidirectional filtering sustains continuous plant–microbe chemical crosstalk and maintains the structural stability of soil microbial communities [5].

Nicotiana tabacum L., an allotetraploid Solanaceous crop with robust root adaptability, is a vital economic crop widely cultivated in China [6,7]. However, intensive cultivation and high nutrient requirements render tobacco prone to soil-borne pathogen infection. Long-term continuous monocropping further weakens plant defensive potential and induces considerable yield losses. As a major flue-cured tobacco production base, Fujian Province features a warm and humid subtropical climate that exacerbates monocropping-related obstacles and triggers frequent outbreaks of tobacco bacterial wilt caused by *Ralstonia solanacearum* [8]. Existing mechanistic studies have verified that rhizosphere microbial dysbiosis disrupts plant immune homeostasis and lowers host tolerance to prevalent soil-borne pathogens, including *Rhizoctonia solani*, *R. solanacearum*, and *Fusarium oxysporum* [9]. Microbial community imbalance disrupts rhizosphere species interaction patterns, reduces community diversity, and creates favorable niche conditions for opportunistic pathogens. Meanwhile, impaired recognition of microbe-associated molecular patterns (MAMPs) compromises plant immune surveillance and further attenuates host disease resistance, as documented in previous studies [10,11]. Consistent with published findings, healthy tobacco rhizospheres actively recruit beneficial microbial taxa, primarily Aspergillus spp., which are negatively correlated with the abundance of Fusarium pathogens. In contrast, diseased rhizospheres harbor lower microbial diversity and vacant ecological niches, which facilitate the colonization and proliferation of pathogenic microorganisms [12].

Conventional management of tobacco soil-borne diseases predominantly depends on synthetic chemical pesticides, which account for approximately 15% of global agricultural fungicide consumption each year [13]. Despite their efficient suppression of pathogen proliferation and disease progression, chemical pesticides often destabilize indigenous rhizosphere microbial assemblages, impair plant immune competence, and induce long-term ecotoxicological risks in agricultural soils [14]. Microbial biocontrol agents have emerged as sustainable alternatives for soil-borne disease management. Nevertheless, the complexity of rhizosphere microhabitats and the poor field colonization capacity of inoculated microbes substantially restrict their practical application efficacy, with the field biocontrol efficiency against tobacco bacterial wilt generally lower than 60% [15,16]. In this context, exploring innovative, sustainable strategies is essential to break through the existing bottlenecks of current disease management regimes.

PQQ is a ubiquitous endogenous redox cofactor that participates in biological energy metabolism and electron transport chain reactions across diverse organisms [17]. PQQ improves microbial environmental adaptability and stress resistance by eliminating reactive oxygen species and modulating stress-responsive gene expression [18]. Distinct from traditional chemical pesticides, PQQ serves as a natural biological biostimulant. Low-dose PQQ supplementation facilitates plant growth and remodels host energy metabolism and hormone signaling networks [19,20]. Although the physiological functions of PQQ have been partially documented, how exogenous PQQ mediates the assembly and functional differentiation of tobacco rhizosphere microbial communities remains largely uncharacterized. In particular, systematic metagenomic evidence clarifying its dose-dependent regulatory patterns in tobacco rhizosphere microbiota is still lacking.

Focusing on tobacco rhizosphere soil, this study adopted metagenomic sequencing to decipher the compositional variation, diversity dynamics, and functional profiling of microbial communities under gradient PQQ treatments. We hypothesized that PQQ modulates tobacco rhizosphere microbial communities in a typical concentration-dependent manner, and optimal PQQ dosage could restructure microbial community assembly toward a structurally balanced microbiota. The present study focused on characterizing microbial community responses to graded PQQ supplementation, aiming to clarify the PQQ-driven reshaping patterns of tobacco rhizosphere microbiota. This work provides fundamental theoretical insights and technical references for the development of sustainable soil-borne disease management strategies in tobacco cultivation.

## 2. Materials and Methods

### 2.1 Experimental Design and Plant Materials

PQQ, (98% HPLC) was purchased from Aladdin Biochemical Technology Co., Ltd. (Shanghai, China; Cat# D134423). The field trial was conducted in 2026 at a tobacco planting base located in Xiacun Village, Nakou Town, Shaowu City, Nanping City, Fujian Province, China (Longitude: 117°38′50.09″E, Latitude: 27°08′21.39″N). The experiment adopted a soil drenching method to investigate the concentration-gradient effects of exogenous PQQ on rhizosphere microbiome assembly and function. Rhizosphere soil samples were collected from uniformly growing tobacco plants in the field for subsequent in-situ field experiments. The experiment adopted a soil drenching method to investigate the concentration-gradient effects of exogenous PQQ on rhizosphere microbiome assembly and function. Four treatments were established: a water control (CK) and three PQQ concentrations (0.5 μmol/L [A], 1 μmol/L [B], and 2 μmol/L [C]). Each treatment was set up with three biological replicates in a randomized complete block design. Each replicate plot, with an area of 199.83 m², contained a total of 88 tobacco plants with unified planting specifications. Throughout the experimental period, only water and PQQ solutions were applied via soil drenching, while all other field management practices remained consistent across all plots. Throughout the entire growth experimental period, only sterile water and gradient PQQ solutions were applied via root irrigation; all other cultivation practices (weeding, watering, pest exclusion without chemical fungicides) remained consistent across all plots to avoid cross-interference.

### 2.2 Soil Sampling and Preservation

Rhizosphere soil sampling followed the standardized shake-off rhizosphere separation protocol: tobacco roots were gently shaken to remove loosely attached bulk soil, and fine soil particles tightly adhered to root epidermal surfaces (0-20 cm topsoil layer) were collected as rhizosphere samples. Three biological replicates were collected for each treatment treatment. All soil samples were sieved through a 2 mm stainless steel mesh to remove root debris, stone particles and plant residues. Subsequently, qualified soil samples were rapidly frozen in liquid nitrogen, and long-term preserved at -80 °C ultra-low temperature refrigerator for subsequent metagenomic library construction and sequencing analysis.

### 2.3 Metagenomic Library Construction and Sequencing

Genomic DNA was extracted from rhizosphere soil using the BGI Optimal series universal DNA library preparation kit (Catalog LD00R08). DNA quality control thresholds were defined as: DNA concentration ≥12.5 μg/μL, fragment length >20 kb, free of protein, RNA and salt contamination. Only qualified DNA templates were used for subsequent library preparation. Genomic DNA was randomly fragmented by Covaris ultrasonication system, and 200-400 bp DNA fragments were enriched and purified using magnetic bead separation. The library was constructed following standard protocols: end-repair, 3′ A-tailing, and adapter ligation. The ligated products were amplified by PCR, purified using magnetic beads, and quantified. The purified libraries were denatured to generate single-stranded circular DNA templates. After removing linear DNA, the qualified libraries were sequenced on the MGISEQ-2000 platform using paired-end 150 bp (PE150) chemistry.

### 2.4 Taxonomic Profiling and Annotation

Raw sequencing data were quality-filtered using SOAPnuke v2.3[21]. Host-derived reads were aligned and filtered out using Bowtie2 v2.4.4[22] based on the published tobacco reference genome (Nicotiana tabacum, GCF_000715135.1), and high-quality clean data were obtained for subsequent bioinformatic analysis. Taxonomic classification of clean reads was performed using Kraken2 v2.1.2[23]. This tool decomposed sequencing reads into *k*-mers and assigned taxonomic labels by querying against the NCBI nt database. The Lowest Common Ancestor (LCA) algorithm was employed to assign classifications across all taxonomic ranks (Kingdom, Phylum, Class, Order, Family, Genus, Species). To correct for potential biases in abundance estimates caused by read mapping ambiguity, Bracken v2.6.1[24] was utilized to re-estimate species abundances based on a Bayesian probability model which read length parameter set to 150 bp. Based on the corrected normalized abundance matrices, taxonomic information at the kingdom, phylum, and genus levels were extracted. The relative abundances of bacteria, archaea, fungi, viruses, and unclassified treatments were statistically analyzed to characterize the structure and composition of the rhizosphere soil microbiome.

### 2.5 Functional Profiling and Annotation

*De novo*assembly of high-quality reads was performed using MEGAHIT v1.2.9[25] based on a *k*-mer algorithm to generate contiguous sequences (contigs). Contigs shorter than 300 bp were discarded. Assembly continuity and completeness were evaluated using metrics including N50, N90, and total sequence length. Open reading frames (ORFs) were predicted from the assembled contigs using MetaGeneMark v3.38[26] to obtain the full gene sequences. Redundancy removal was conducted using CD-HIT v4.8.1[27] to construct a non-redundant gene catalog. Gene abundance quantification was performed using Salmon v1.6.0[28], generating a relative abundance matrix. Functional annotation of the non-redundant genes was carried out using the blastp module of DIAMOND v0.8.24[29]. To correct for biases arising from gene length and sequencing depth, Transcripts Per Million (TPM) normalization was applied. Annotated genes were aligned against major functional databases, including KEGG v101.0[30], COG 20201125[31], eggNOG eggnog 5.0[32], Swiss-prot release-2021 04[33], BacMet2 20180311[34],CAZy 20240326[35], and. Additionally, antibiotic resistance genes (ARGs) were annotated using RGI v5.2.1 with default parameters based on the CARD v3.0.9 database[36].

Transcripts Per Million:

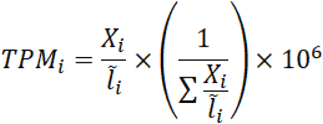

Transcripts Per Million: where i= : Gene index; *X_i_* = read counts mapped to gene i; *Ĩ_i_* = average length of gene i.

### 2.6 Statistical Analysis and Visualization

Statistical analyses were performed using Microsoft Excel. Alpha diversity indices including Shannon, Simpson, Chao1 were calculated using QIIME2. Beta diversity was assessed based on Bray-Curtis dissimilarity and visualized by Principal Coordinates Analysis (PCoA). Differential abundance of functional pathways was analyzed using STAMP with Benjamini-Hochberg FDR correction (*p* < 0.05). All experimental data are presented as means ± standard deviation (Mean ± SD). Figures were generated with Origin Pro 2025. Multi-omics downstream visualization of metagenomic datasets was performed on Dr. Tom cloud platform (https://biosys.bgi.com).

## 3. Result

All metagenomic datasets, taxonomic profiles and functional annotations were systematically analyzed to reveal concentration-dependent reshaping patterns of tobacco rhizosphere microbiome induced by exogenous PQQ gradient treatments. The overall workflow contained assembly quality evaluation, taxonomic diversity analysis, community composition characterization, differential biomarker screening and multi-database functional profiling.

### 3.1 *De novo* Assembly and Gene Prediction

Prior to taxonomic and functional annotation, de novo metagenomic assembly was performed on quality-filtered reads from rhizosphere soil samples of four experimental treatments. Comprehensive assembly metrics, including total contig number, N50, N90, and average contig length, are summarized in Table 1-1. The N50 values ranged from 562 to 736 bp, while N90 values varied between 347 and 388 bp, with average contig lengths of 576 to 711 bp. These assembly statistics indicate adequate sequence continuity for downstream open reading frame (ORF) prediction and functional gene catalog construction.

**Table 1-1.**
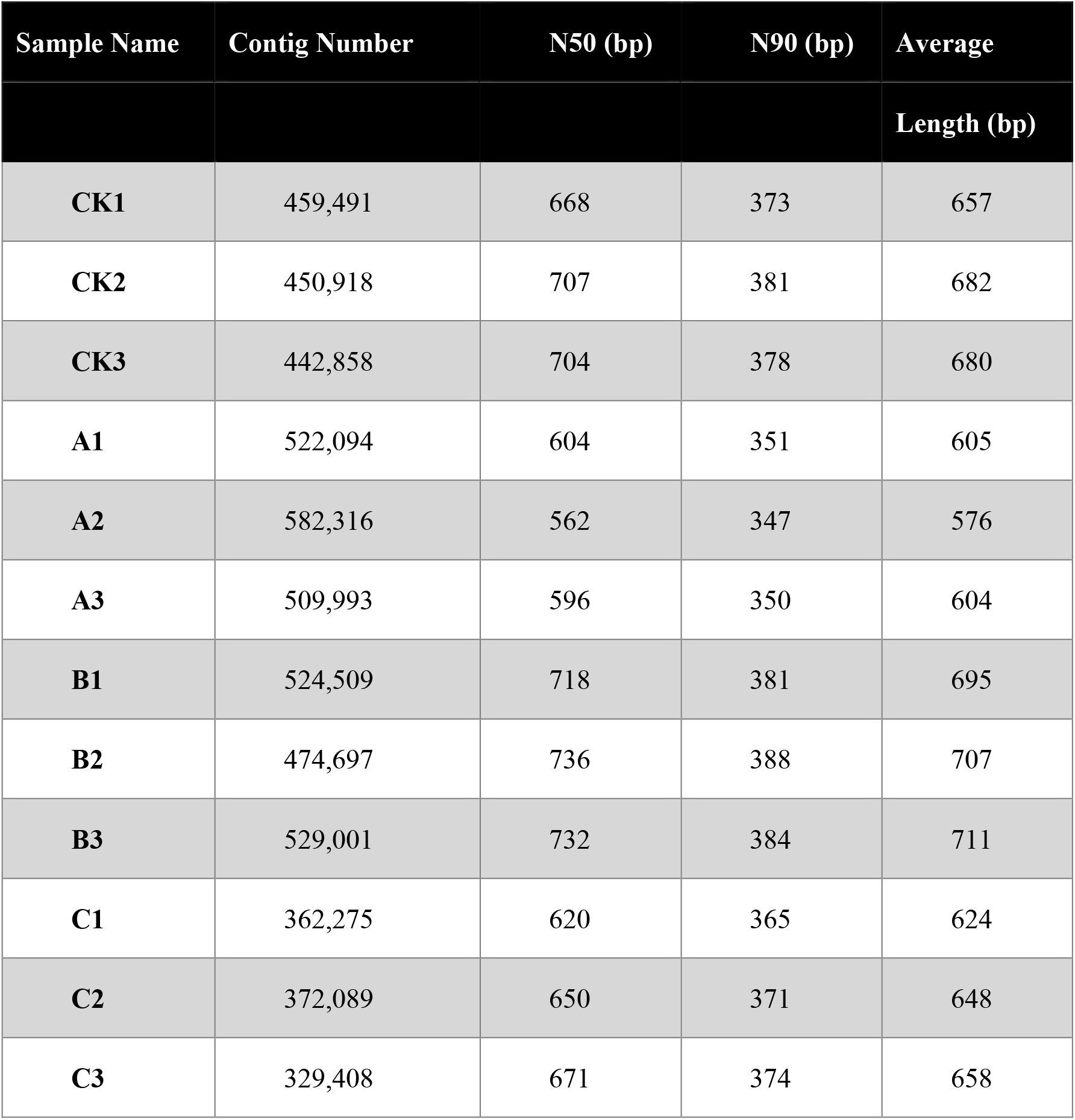
Summary statistics of metagenomic assembly for CK and PQQ-treated treatments.

Global gene prediction was conducted on assembled contigs. The number of predicted coding sequences (CDS) per sample ranged from 474,862 to 808,636 (Table 1-2). Notable inter-treatment differences were observed: the low-dose (A) and moderate-dose (B) PQQ treatments generated substantially more raw CDS than the control (CK), whereas the high-concentration treatment (C) produced the fewest CDS among all treatments. Redundancy removal was performed at a threshold of 95% sequence identity and 90% coverage to eliminate homologous and repetitive sequences. Consequently, the final non-redundant gene catalog was constructed, and per-sample gene abundances ranged from 430,437 to 731,973 TPM-mapped genes.

**Table 1-2.**
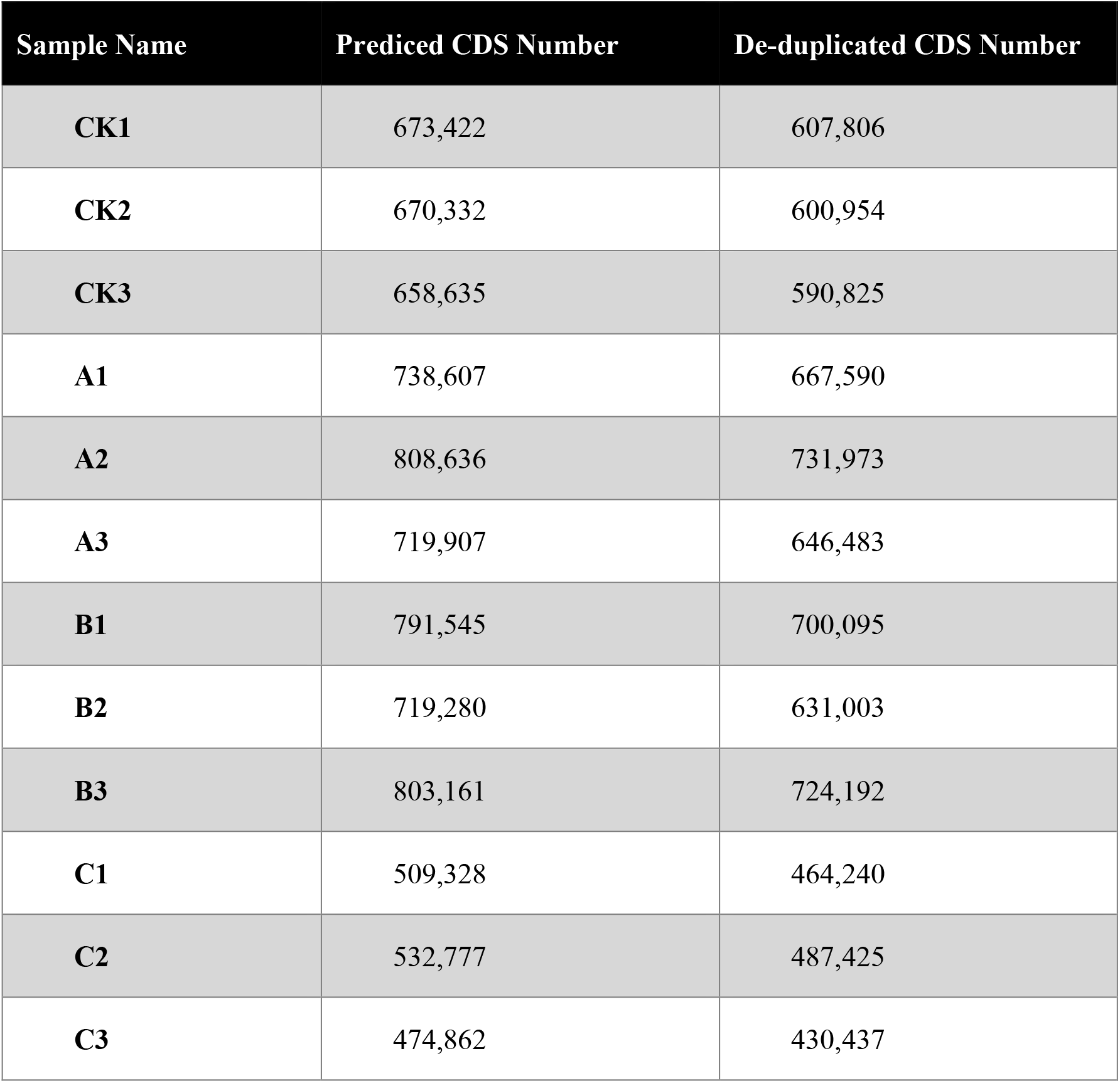
Statistics of gene prediction and non-redundant gene catalog construction.

### 3.2 Taxonomic Diversity

Alpha and beta diversity indices were jointly calculated to quantify how PQQ dosage modulated the overall richness, uniformity and structural separation of tobacco rhizosphere microbial communities.

#### 3.2.1 Alpha Diversity Indices

Three core αlpha diversity indices, including Shannon, Simpson and Chao1, were used to assess the evenness and species richness of tobacco rhizosphere microbial communities under gradient PQQ treatments (**Table 2**). Microbial community evenness displayed a distinct U-shaped trend along the PQQ concentration gradient. The Shannon and Simpson indices showed no significant differences among the CK, low-dose (A) and high-dose (C) treatments but decreased significantly in the moderate-dose (B) treatment. The Chao1 index, which reflects microbial species richness, exhibited varied responses to PQQ supplementation. Both the A and B treatments produced significantly higher Chao1 values than CK, while no significant difference in species richness was observed between the CK and C treatments. One-way ANOVA further confirmed that the 1 μmol/L PQQ treatment (B) significantly reduced community evenness relative to all other treatments. Collectively, these alpha diversity results indicated that the moderate PQQ dosage (1 μmol/L) exerted the most prominent modulatory effect on the assembly of tobacco rhizosphere microbial communities. **Table 2** Alpha diversity indices of rhizosphere soil microbial communities under different treatments.

**Table 2.**
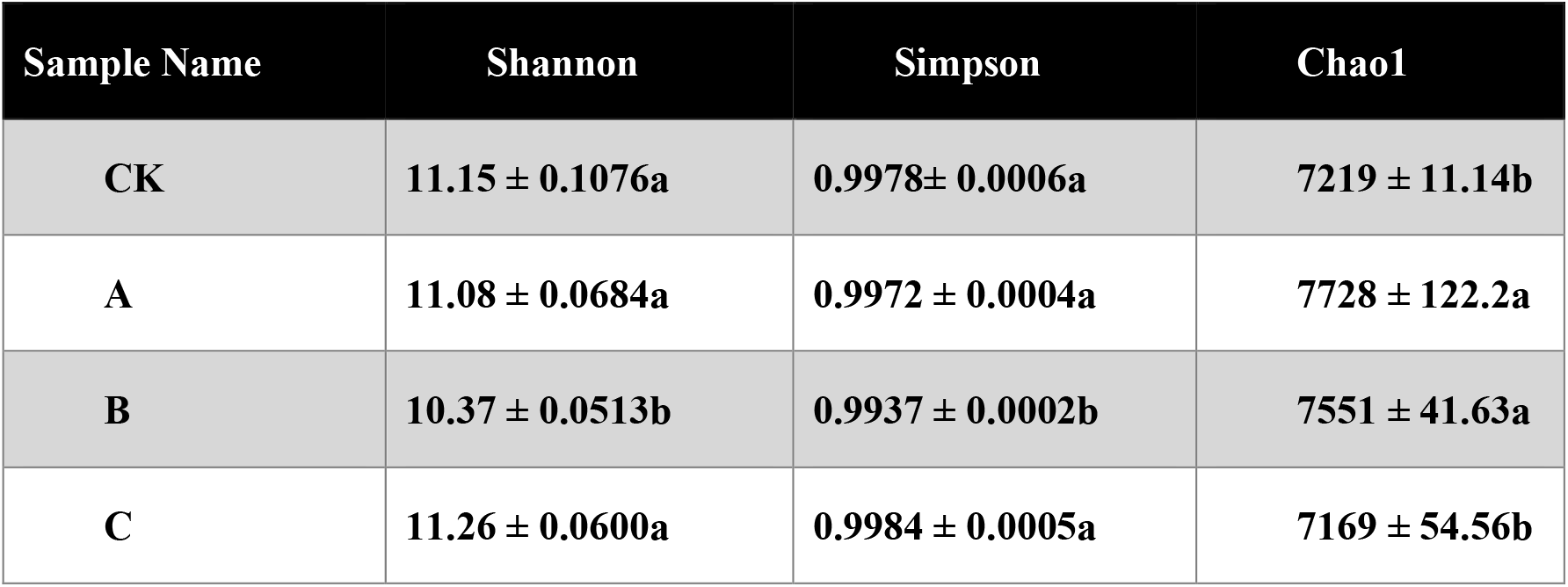
Alpha diversity indices of rhizosphere soil microbial communities under different treatments.

#### 3.2.2 Beta Diversity and Community Structure

To investigate the structural variations in the rhizosphere microbiome, we employed Principal Coordinates Analysis (PCoA), Non-metric Multidimensional Scaling (NMDS), and Analysis of Similarities (ANOSIM) based on the Bray-Curtis distance matrix. PCoA (**Figure 1A**) explained 72.43% of the total variance on the first axis (PCoA1) and 13.67% on the second axis (PCoA2). The NMDS ordination plot (**Figure 1B**) showed no obvious separation between the low-dose PQQ treatment (A) and the CK treatment, while the moderate and high PQQ concentration treatments (B and C) exhibited distinct spatial segregation from CK. Samples within each treatment clustered densely, with pronounced disparities observed among different treatments. NMDS yielded a stress value of 0.0173. Treatment B was clearly separated from CK, A, and C. Furthermore, ANOSIM (**Figure 1C**) revealed an R-value of 1.0 (*p*=0.001), showing that inter-treatment variation was substantially larger than the intra-treatment variation among microbial communities.

**Figure 1.**
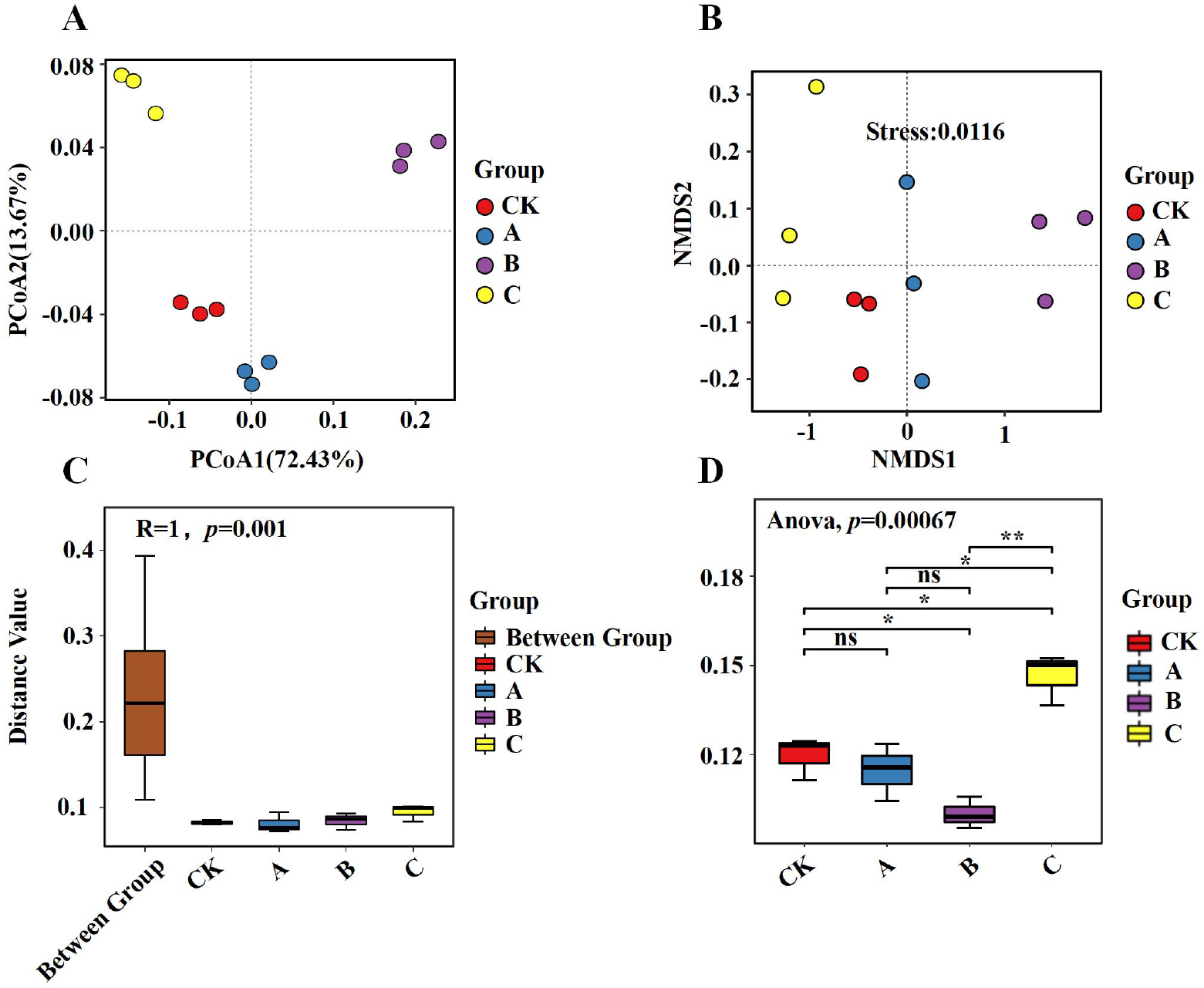
Effects of varying concentrations of PQQ on the structure of the tobacco rhizosphere bacterial community. (**A**)PCoA based on Bray-Curtis dissimilarity. (**B**)NMDS ordination (stress = 0.0116) showing the separation of bacterial communities among treatments. (**C**)ANOSIM confirming significant differences in community composition between Treatment B and other treatments (R=1, *p*<0.05). (**D**)Boxplot illustrates the distribution of Bray-Curtis beta diversity distances within intra-treatment and between inter-treatment.

To further quantify the dissimilarity in microbial community structure across treatments, a boxplot was generated based on the Jensen-Shannon Divergence (JSD) distance matrix (**Figure 1D**). The inter-treatment distances between the PQQ-treated treatments and the CK treatment were significantly higher than the intra-treatment distances (ANOVA, *p*=0.00067). Moreover, the compact distribution of the intra-treatment distance boxes indicated high reproducibility and stability among the biological replicates within each treatment.

### 3.3 Taxonomic Composition of the Rhizosphere Microbiome

#### 3.3.1 Relative Abundance of Dominant Taxa

At the Kingdom level, the rhizosphere microbial community was taxonomically classified into four major treatments: Bacteria, Archaea, Eukaryotes, and Viruses (**Table 3**). Bacteria constituted the dominant kingdom across all samples, with relative abundances consistently exceeding 97%. Notably, the B treatment (1 μmol/L PQQ) harbored the highest bacterial abundance, surpassing 99%. Archaea represented the second most abundant treatment, accounting for 0.65%-1.84% of the total sequences. Eukaryotes and Viruses were present as minor components, comprising 0.19%-0.26% and 0.01%-0.04%, respectively. Although the community was overwhelmingly dominated by Bacteria, the rhizosphere sustains a complex microbiome.

**Table 3.**
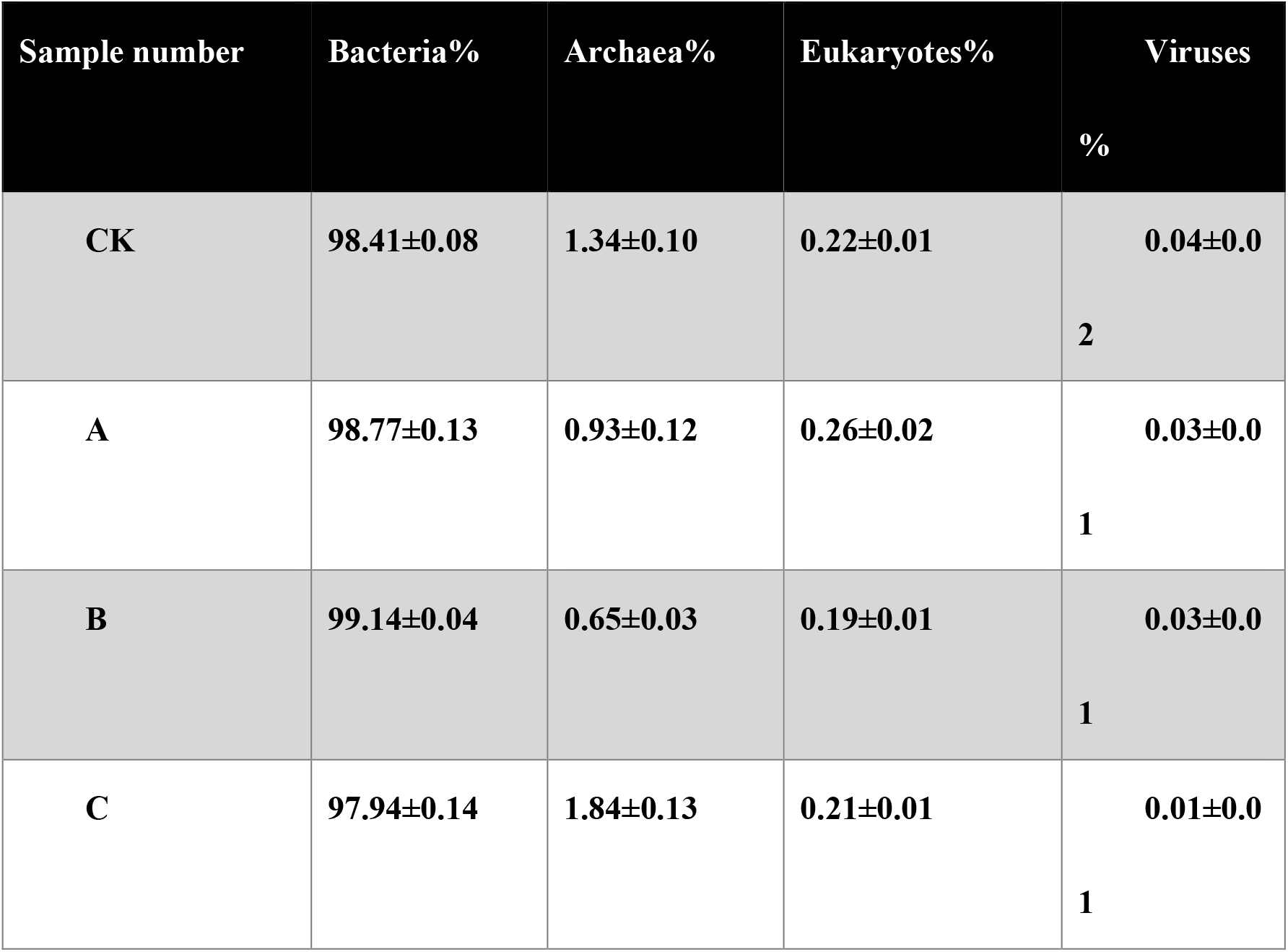
Relative abundance of rhizosphere microbial communities at the Kingdom level.

#### 3.3.2 Taxonomic Distribution at Phylum and Genus Levels

Variations in rhizosphere microbial taxonomic structure were explored at the phylum level by analyzing the top 20 dominant bacterial taxa across all experimental treatments (**Figure 2A**). The rhizosphere microbial community was predominated by Pseudomonadota and Actinomycetota, whose cumulative relative abundance exceeded 70% across all samples, serving as the primary phyla constituting local microbial assemblages. In the CK treatment, Pseudomonadota and Actinomycetota accounted for approximately 60% and 25% of total microbial sequences, respectively. Gradient PQQ supplementation reshaped the abundance profile of these dominant phyla. Treatment B (1 μmol/L) harbored the highest proportion of Pseudomonadota, reaching 66%. By comparison, the abundance of Pseudomonadota in low-dose (A, 0.5 μmol/L) and high-dose (C, 2 μmol/L) treatments was comparable to control levels. The relative proportion of Actinomycetota decreased progressively with elevated PQQ dosage, recording 29% in treatment A, 24% in treatment B and 22% in treatment C.

**Figure 2.**
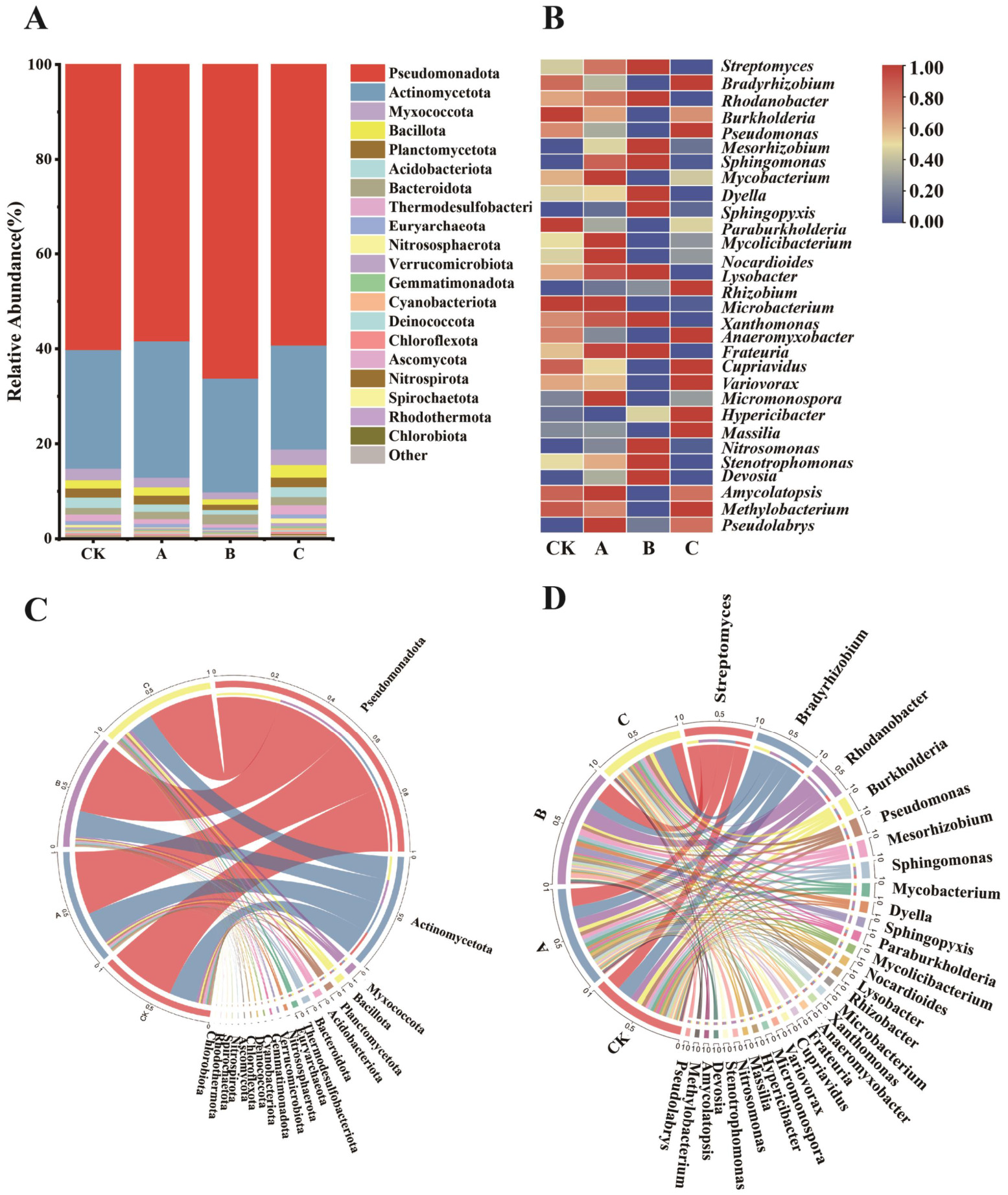
Taxonomic composition and Circos visualization of the tobacco rhizosphere bacterial community under different PQQ treatments. **(A)** Stacked bar chart showing the relative abundance of the top 20 dominant bacterial phyla across all samples. (B) Heatmap displaying the relative abundance of the top 30 dominant bacterial genera (remaining genera pooled as “Others”) (C) Circos plot displaying the distribution and proportion of dominant bacterial phyla among treatments. (D) Circos plot illustrating the distribution and proportional contribution of dominant bacterial genera among treatments.

At the genus level, the top 30 abundant microbial genera were subjected to hierarchical clustering analysis and visualized via heatmap (**Figure 2B**). The control rhizosphere microbiota was primarily composed of*Burkholderia*, *Paraburkholderia*, and *Microbacterium*. PQQ amendment altered genus-level community composition, with noticeable accumulation of *Streptomyces*, *Rhodanobacter*, *Sphingomonas*, *Lysobacter*, and *Frateuria*. Several actinobacterial taxa, including *Mycobacterium*, *Mycolicibacterium*, *Nocardioides*, and *Micromonospora*, were enriched under low-concentration PQQ treatment but declined gradually with increasing PQQ dosage. Multiple beneficial genera, including *Rhodanobacter*, *Mesorhizobium*, *Sphingomonas*, *Devosia*, *Lysobacter*, *Nitrosomonas*, *Stenotrophomonas*, and *Dyella*, accumulated from CK to low and moderate PQQ treatments and declined sharply under high-concentration treatment (C). By contrast, *Bradyrhizobium*, *Pseudomonas*, *Hypericibacter*, and *Massilia* exhibited reduced abundance at low PQQ dosage and recovered prominently at the highest concentration. Overall, microbial taxa displayed dosage-dependent variation in abundance in response to gradient PQQ treatment.

#### 3.3.3 Integrative Analysis of Taxonomic Composition via Circos Plot

Complementary Circos plot analysis further visualized the correspondence between experimental treatments and rhizosphere microbial profiles at both the phylum and genus levels, illustrating the relative proportions of taxa and their distribution across treatments. At the phylum level (**Figure 2C**), Pseudomonadota and Actinomycetota dominated all samples; their combined relative abundance far exceeded that of the remaining taxa, identifying them as the core components of the tobacco rhizosphere microbiome. At the genus level (**Figure 2D**), *Streptomyces*, *Bradyrhizobium*, *Rhodanobacter*, *Burkholderia* and *Pseudomonas* were the five most abundant genera. Each genus exhibited a distinct distribution pattern across the control and PQQ treatments, with abundance shifting non-linearly along the PQQ concentration gradient.

#### 3.3.4 Differential Biomarker Taxa Identified by LEfSe Analysis

Given that treatment B produced the most pronounced shifts in αlpha and βeta diversity, Linear Discriminant Analysis Effect Size (LEfSe) was applied to identify the taxa that discriminated treatment B from the CK treatment (LDA score > 2.0, *p* < 0.05) in **Figures 3A** and **3B**. Both the cladogram and the corresponding LDA score plot revealed clear taxonomic divergence between the two treatments, with several genera identified as diagnostic biomarkers. *Streptomyces*, *Sphingomonas*, *Rhodanobacter* and *Dyella* were specifically enriched in treatment B, with LDA scores exceeding 3.0. conversely, *Pseudomonas*, *Burkholderia* and *Mycobacterium* were enriched in the CK treatment (LDA < -3.0). These distinct biomarker distributions highlighted consistent compositional differences between the moderate PQQ treatment and the control rhizosphere microbiome.

**Figure 3.**
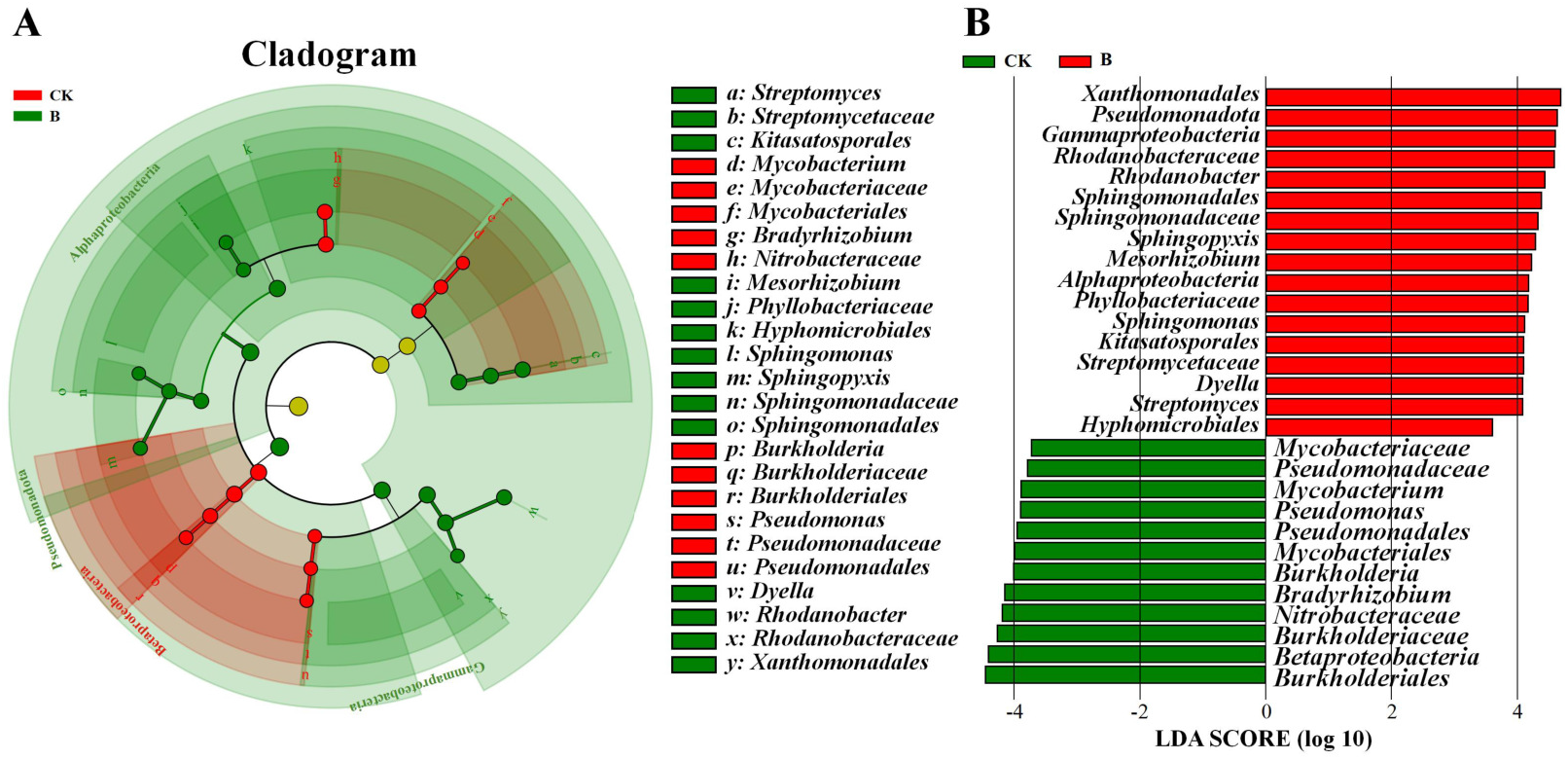
LEfSe identifying biomarker bacterial taxa between Treatment B and the CK. (**A**) Cladogram illustrates the phylogenetic distribution of bacterial taxa. Nodes are colored according to their enrichment in Treatment B (green) or CK (red). (**B**) Histogram of LDA scores. Positive scores indicate enrichment in Treatment B, whereas negative scores indicate enrichment in CK.

### 3.4 Functional Annotation and Differential Analysis

#### 3.4.1 KEGG Functional Profiling and Differential Analysis

To investigate the functional shifts in the rhizosphere microbiome following PQQ application, the top 20 most abundant Level 2 KEGG pathways were selected for visualization (Figure 4A). The functional landscape was dominated by Global and overview maps, Carbohydrate metabolism, Amino acid metabolism, Energy metabolism, and Metabolism of cofactors and vitamins.

**Figure 4.**
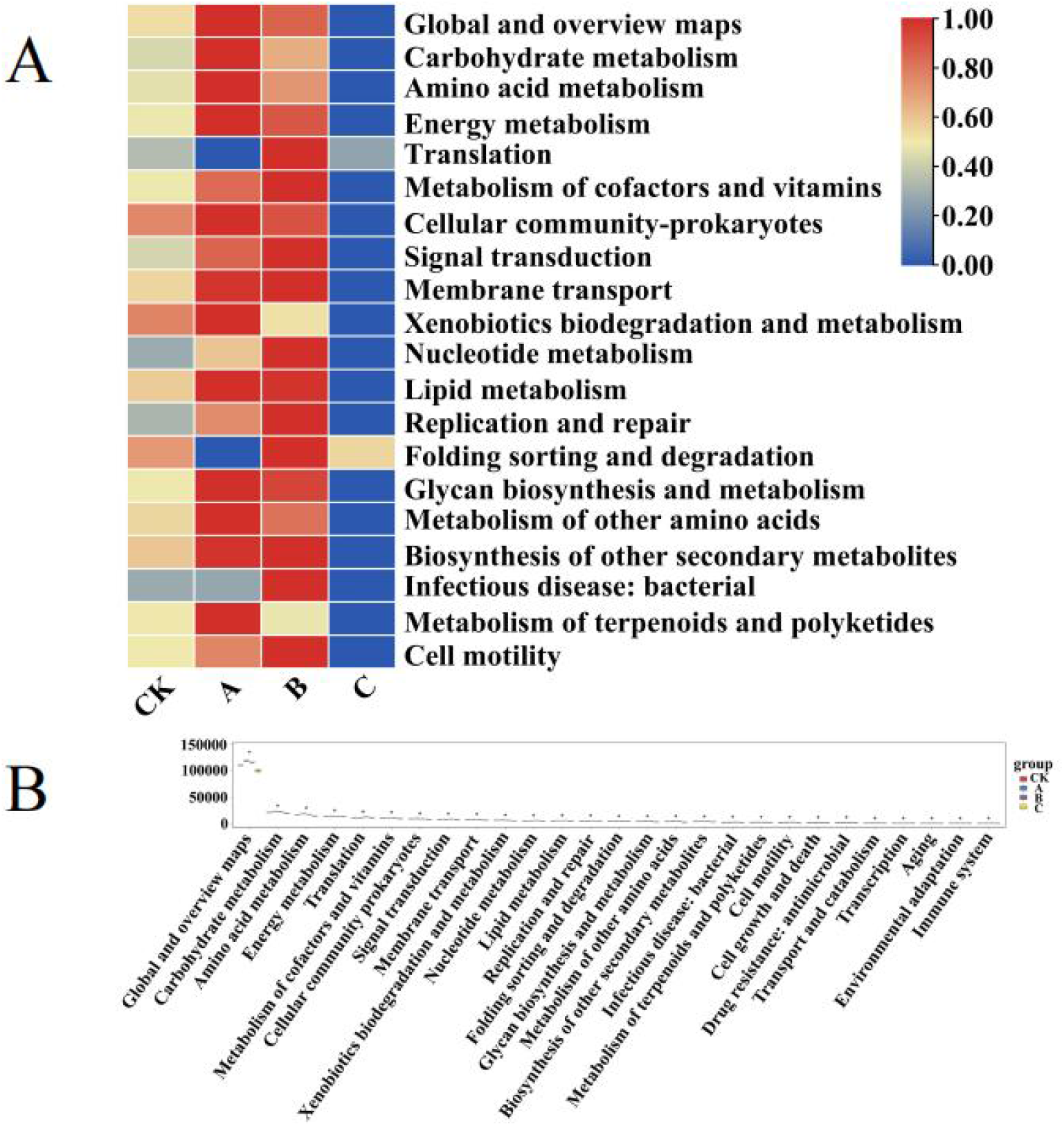
Differential abundance of KEGG Level 2 functional pathways under different PQQ treatments. (A) Heatmap showing the relative abundance of KEGG Level 2 pathways across all samples. (B) Boxplots comparing the abundance of selected KEGG Level 2 pathways between Treatment A、B、C and CK. Statistical significance was determined using Kruskal-Willas (∗*p* < 0.05, ∗∗*p* < 0.01).

Overall, the functional profiles exhibited a non-monotonic dose-response to PQQ exposure. The majority of metabolic pathways, including Global and overview maps, Carbohydrate metabolism, Amino acid metabolism, Energy metabolism, Cellular community-prokaryotes, Signal transduction, and Membrane transport, showed elevated abundances in Treatments A and B compared with CK, but were markedly suppressed in Treatment C. Notably, Translation, Nucleotide metabolism, Folding sorting and degradation, Infectious disease: bacterial, and Cell motility peaked specifically under Treatment B (1 μmol/L PQQ), exhibiting lower levels in both CK and Treatment A. Conversely, Xenobiotics biodegradation and metabolism and Metabolism of terpenoids and polyketides remained relatively stable across CK, A, and B, but declined under Treatment C. These results indicate that low to moderate PQQ concentrations enhance the functional potential of the rhizosphere microbiome, whereas high concentration exerts a broad suppressive effect..

To further elucidate the regulatory effects of PQQ on the metabolic potential of the rhizosphere microbiome, abundance variations of KEGG Level 2 pathways were analyzed using box plots combined with statistical hypothesis testing (Figure 4B). Kruskal-Wallis tests revealed significant inter-treatment differences across all 27 KEGG Level 2 pathways (FDR < 0.05), indicating a systematic restructuring of the functional repertoire in response to PQQ treatment.

Global and overview maps represented the most abundant functional category across all treatments, followed by core metabolic modules including Carbohydrate metabolism, Amino acid metabolism, and Energy metabolism. Notably, the majority of pathways exhibited elevated abundance in Treatment B (1 μmol/L PQQ), which was significantly higher than that in CK and other treatment treatments. In sharp contrast, Treatment C (2 μmol/L PQQ) showed markedly suppressed abundance across nearly all pathways, with levels approaching or falling below those of the CK treatment. Collectively, these findings demonstrate that moderate PQQ application significantly enhances the metabolic network architecture of the rhizosphere microbiome, whereas excessive PQQ exerts a broad suppressive effect.

#### 3.4.2 COG Distribution and Intertreatment Differential Analysis

Functional annotation against the COG database identified 25 functional categories in Figure 5. The dominant classes included Translation, ribosomal structure and biogenesis, General function prediction only, Energy production and conversion, Transcription, Amino acid transport and metabolism, Signal transduction mechanisms, Posttranslational modification, protein turnover, chaperones, Cell wall/membrane/envelope biogenesis, and Coenzyme transport and metabolism.

**Figure 5.**
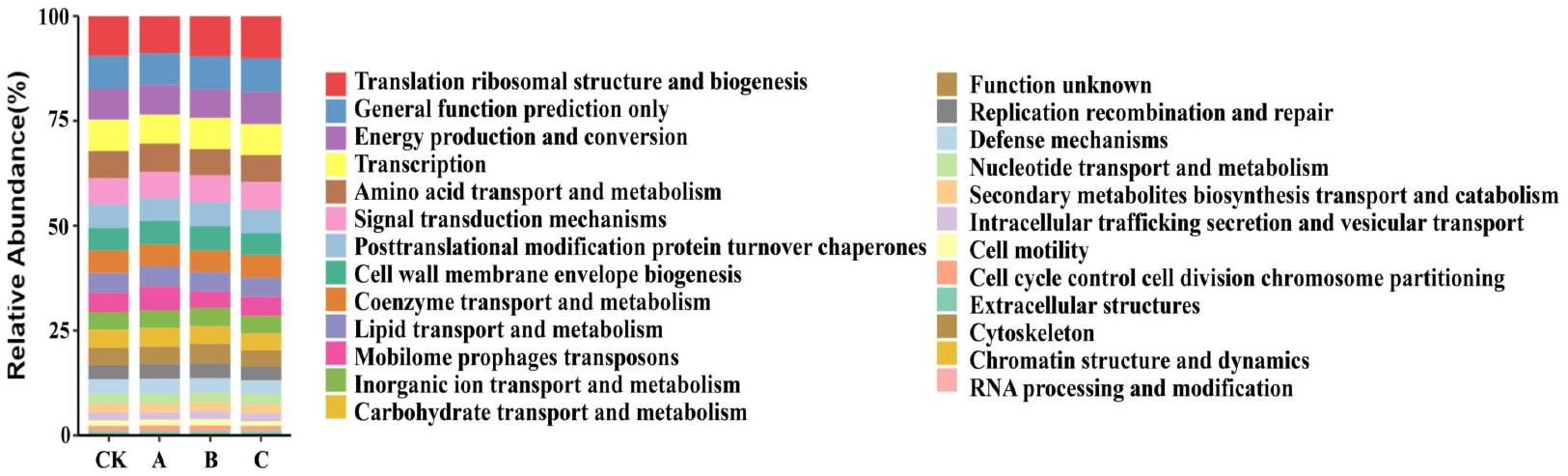
Distribution of COG functional categories in the tobacco rhizosphere microbiome under different PQQ treatments.

STAMP analysis revealed significant differences in 21 of the 25 annotated COG categories across treatments (FDR *p* < 0.05) in Figure 6. Treatment B (1 μmol/L PQQ) exhibited the highest mean gene abundance across most functional categories, while the CK treatment showed the lowest levels; treatments A and C displayed intermediate abundance. The most distinct inter-treatment variations were detected in transcription (K), amino acid transport and metabolism (E), posttranslational modification, protein turnover, and chaperones (O), cell wall/membrane/envelope biogenesis (M), and general function prediction only (R).

**Figure 6.**
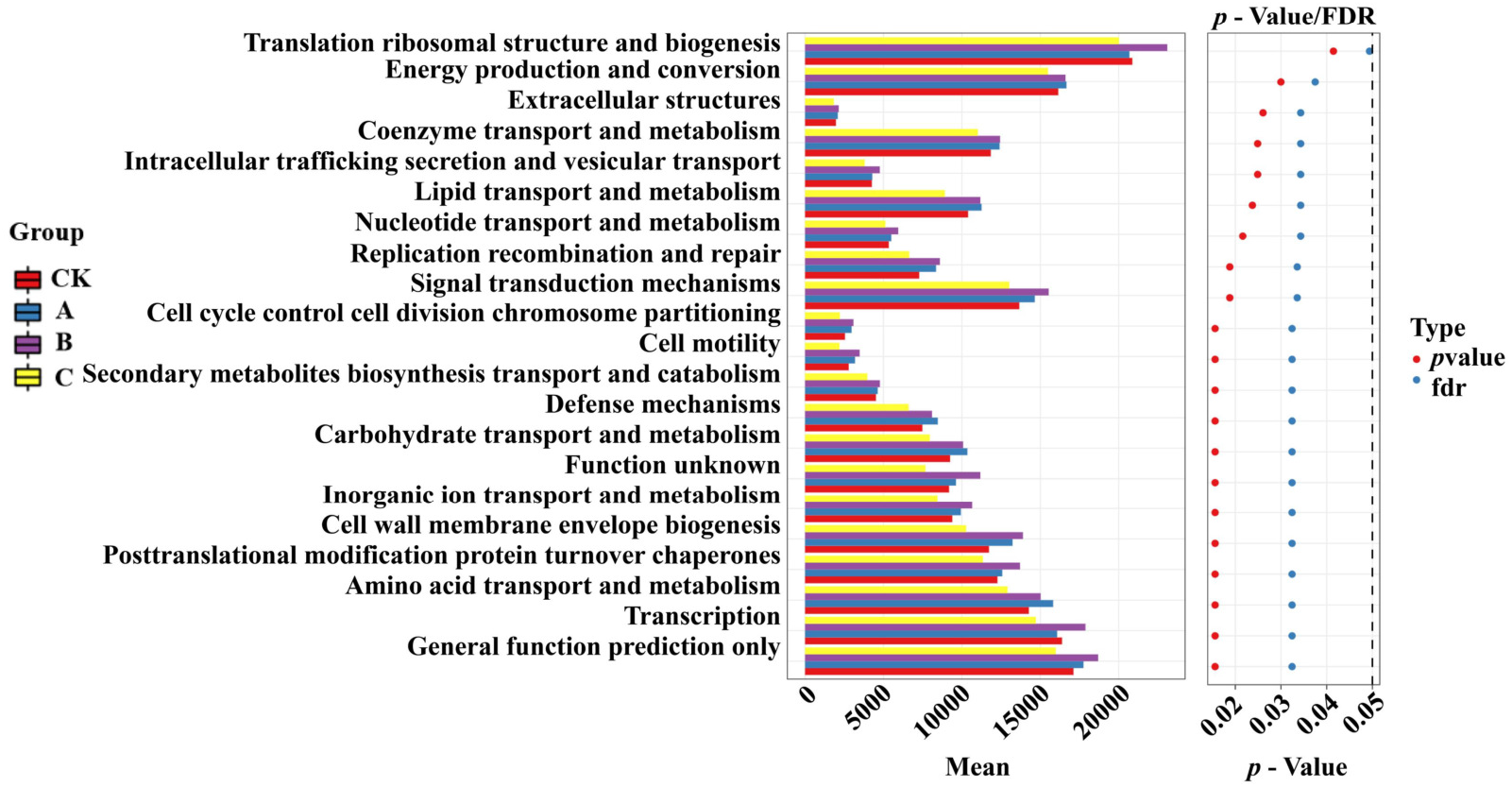
Differentially abundant Clusters of Orthologous Treatments (COG) functional categories between Treatment A、B、C and CK.

#### 3.4.3 Antibiotic Resistance Gene Analysis

ARG profiles were annotated against version 3.0.9 of the CARD database, and the 30 most abundant categories were retained for analysis, with all remaining low-abundance categories pooled as Others as shown in Figure 7A. Seven categories exceeded 5% relative abundance in all four treatments, namely penam, cephalosporin, carbapenem, tetracycline, macrolide, fluoroquinolone and phenicol. Within this dominant set, beta-lactam resistance genes were the most prominent, with penam, cephalosporin and carbapenem together accounting for approximately 30% of the total ARG relative abundance, whereas the remaining beta-lactam subclasses including cephamycin, monobactam and penem stayed at much lower levels. Two further categories showed treatment-specific predominance: diaminopyrimidine exceeded 5% only in treatment B, while aminoglycoside exceeded 5% in both treatments B and C. All other categories, including lincosamide, streptogramin, peptide, glycopeptide and pleuromutilin, remained below 5% in every treatment.

**Figure. 7.**
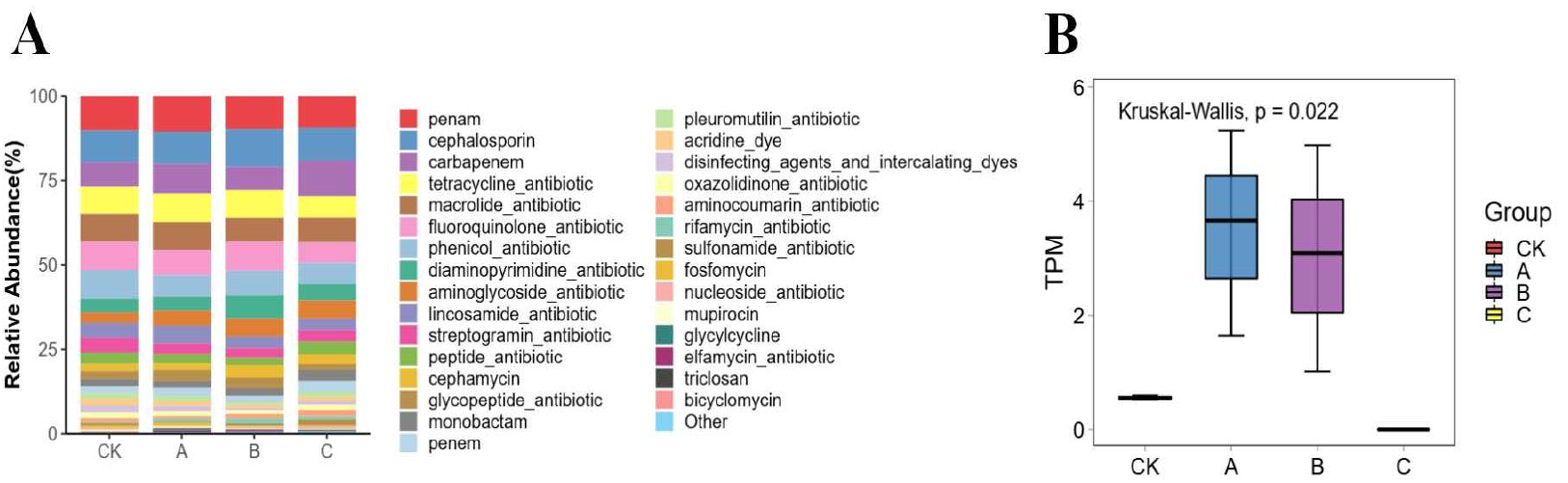
Composition and abundance of the soil resistome across treatment treatments. (A) Relative abundance of the top 30 ARG categories. (B) Comparison of total ARG abundance (TPM) among treatments. CK, A, B, C, treatment treatments.

After Benjamini-Hochberg correction for multiple testing, no ARG category remained significantly different in abundance among the CK, A, B and C treatments, with all FDR values exceeding 0.05. The elfamycin category showed the largest nominal difference, with an uncorrected Kruskal-Wallis *p* value of 0.022 as shown in Figure 7B, but this did not survive correction with an FDR of approximately 0.5.

## 4. Discussion

The rhizosphere microbiome suppresses soil-borne pathogens through nutrient competition, antagonistic compounds and induced host resistance [37]. Root exudates can filter rhizosphere taxa by inducing oxidative stress [38]. PQQ may act in a comparable way. Accordingly, both the taxonomic composition and functional structure of the rhizosphere microbiome differed markedly among the PQQ concentration treatments. However, whether these community-level differences translate into altered soil disease-suppressive capacity remains to be established. Across the PQQ concentration gradient (0.5, 1 and 2 μmol/L), alpha diversity metrics showed marked divergence: species richness, as reflected by the Chao1 index, increased at low and moderate doses and returned to control levels at the highest dose, whereas the Shannon and Simpson indices exhibited a U-shaped response. These patterns align with previous reports of microbial community assembly in agricultural soils [39]. In treatment B, the Shannon and Simpson indices decreased significantly whereas Chao1 richness increased significantly, consistent with a community state of reduced evenness but elevated species richness, in which the relative dominance of a few abundant taxa was enhanced.

Beta diversity analysis showed differences among the PQQ treatments. Principal coordinate analysis separated the four treatments into distinct clusters, and non-metric multidimensional scaling produced a low stress value of 0.0173, indicating that the ordination reliably represented community dissimilarity. Jensen–Shannon divergence was significantly higher between treatments than within treatments at *p* = 0.031, indicating that the compositional variation was attributable to PQQ application rather than to plot-level heterogeneity. These community differences are consistent with previously reported shifts associated with plant-derived redox-active compounds [40,41] and exogenous soil amendments [42]. Along with the alpha diversity profiles, the beta diversity results showed that the rhizosphere microbiome differed among treatments in a concentration-associated, non-monotonic manner. As a redox-active quinone, PQQ may be associated with shifts in rhizosphere redox conditions and thereby favour taxa according to their redox tolerance; however, soil redox potential and metabolite profiles were not quantified, so this possible mechanism remains untested.

At the phylum level, Pseudomonadota and Actinomycetota constituted the core rhizosphere microbiome, participating in nutrient cycling, microbial interactions and plant stress tolerance [43,44], with Actinomycetota being particularly responsive to agricultural management practices [45]. PQQ application was associated with shifts in the relative abundance of several genera within these phyla. *Streptomyces* synthesizes antibiotics and extracellular hydrolases and exhibits biocontrol potential against soil-borne pathogens [46–48]; *Rhodanobacter* functions as a plant growth-promoting endophyte [49], *Sphingomonas* alleviates continuous cropping obstacles via phenolic acid degradation [50], and *Lysobacter* is linked to enhanced soil enzyme activity [51]. These genera all showed higher relative abundance in treatment B. *Xanthomonas*, which influences plant reproductive development [52], and *Frateuria*, a genus reported to harbour a PQQ-dependent alcohol dehydrogenase [53], also displayed treatment-specific variation. Overall, PQQ supplementation was accompanied by substantial compositional shifts, with treatment B showing the most distinct profile characterized by higher relative abundance of taxa functionally linked to nutrient acquisition, antagonism and plant growth promotion. These genera provide promising targets for future culture-dependent validation and rhizosphere microbiome regulation.

KEGG level 2 functional analysis showed higher relative abundance of core metabolic and cellular pathways in treatment B. The overrepresented lipid metabolism pathway is consistent with cell envelope expansion and membrane remodeling during microbial proliferation, a functional trait reported for host-associated symbionts [54]. The higher abundance of signal transduction and cell motility functions has been linked to rhizosphere sensing and chemotactic potential, traits associated with competitive root colonization and pathogen exclusion [55].

COG functional profiling showed a similar pattern. Annotation against the COG database identified 25 functional categories, of which 21 differed significantly among treatments, and treatment B showed the highest mean gene abundance across most categories. This treatment had higher proportions of functions related to translation, ribosomal structure and biogenesis, which are consistent with stable rhizosphere colonization [56], and of energy production and conversion, consistent with sustained metabolic activity [57]. Higher proportions of transcription and of amino acid transport and metabolism have been linked to greater carbon and nitrogen supply for biosynthesis [58,59], whereas higher signal transduction mechanisms have been associated with environmental adaptability [60]. Transcription, amino acid transport and metabolism, cell wall, membrane and envelope biogenesis, and posttranslational modification, protein turnover and chaperones were among the categories showing the largest inter-treatment variation and were also higher in this treatment. Collectively, this treatment was associated with higher relative abundance of functions related to biosynthesis, energy metabolism, colonization and environmental sensing, whereas these functions were lower at the highest concentration.

In addition to the taxonomic and functional profiles, the resistome was examined to assess whether PQQ application was associated with shifts in antibiotic resistance genes. Composition remained broadly stable across the PQQ gradient, with beta-lactam resistance genes predominant throughout. Although ARG abundance showed a non-monotonic numerical pattern, being numerically highest in treatment B and returning to control levels at the highest dose, these differences were not significant after correction for multiple testing. At the concentrations tested, PQQ applied by root irrigation did not show a significant effect on ARG abundance in the tobacco rhizosphere.

In conclusion, the tobacco rhizosphere microbiome responded to PQQ in a concentration-associated, non-monotonic manner, with both taxonomic composition and functional potential showing the most pronounced shifts at 1 μmol/L. Diversity and functional analyses consistently distinguished this treatment, where beneficial genera and functions related to biosynthesis, energy metabolism, colonization and environmental sensing were more abundant. No broad-spectrum antimicrobial effect or significant enrichment of antibiotic resistance genes was detected across the tested concentrations. These findings provide a basis for further evaluation of PQQ in modulating the rhizosphere microbiome in continuous cropping systems.

## 5. Limitations and Future Perspectives

Several limitations should be noted. Metagenomic sequencing captures relative abundance and functional potential, so the differences reported here are compositional rather than quantitative, and their correspondence to microbial activity remains to be tested. Soil physicochemical properties were not determined; however, the treatments were established within a single field under uniform management, and compositional variation was attributable to PQQ application rather than to plot heterogeneity, making baseline soil differences an unlikely explanation. Plant growth and disease incidence were not monitored, so any link between microbiome variation and plant performance remains unconfirmed. Within these constraints, this study demonstrates a clear concentration-associated response of the tobacco rhizosphere microbiome to PQQ, providing a framework for future validation in continuous cropping systems.

## Data availability

The raw sequencing data generated in this study have been deposited in the NCBI Sequence Read Archive (SRA) under BioProject accession PRJNA1462743 (TaxID: 410658). The BioSample accessions are SAMN61519520 to SAMN61519528. The data can be accessed at: https://www.ncbi.nlm.nih.gov/bioproject/PRJNA1462743.

## Acknowledgements

This work was financially supported by Fujian Provincial Natural Science Foundation of China (Grant No. 2026J01310808), Science and Technology Plan Project of Fujian Branch, China Tobacco Corporation (Grant No. KZXH2502B) and National Natural Science Foundation of China (Grant No. 22377012).

